# A novel method for sampling subgingival microbiome - a comparative metatranscriptomic study

**DOI:** 10.1101/2023.02.28.530452

**Authors:** Diana Demusaj, Ryan Toma, Tanveer Khan, Lan Hu, Guruduth Banavar, Momchilo Vuyisich

## Abstract

The subgingival microbiome has been implicated in several oral and systemic diseases, such as periodontitis, arthritis, and Alzheimer’s disease. However, subgingival sampling is challenging and cannot be readily performed outside of primary care facilities. In order to support global, diverse, direct-to-participant clinical research studies, we developed a novel method of sampling the subgingival microbiome by rotationally swabbing the supragingival area, which we named subgingival-P (for proxy) samples. To validate this method, we sampled and metatranscriptomically analyzed subgingival and subgingival-P samples of three different teeth in 20 individuals. The subgingival-P samples were comparable to the subgingival samples in the relative abundances of microorganisms and microbial gene expression levels. Our data demonstrate that the novel method of collecting and analyzing the subgingival-P samples can act as a proxy for the subgingiva, paving the way for large and diverse studies investigating the role of the subgingival microbiome in health and disease.

## Introduction

The microorganisms in the oral cavity are important in maintaining an individual’s oral and systemic health ^1,2^. The oral microbiome comprises viruses, phages, fungi, protozoa, archaea, and bacteria. The bacterial communities alone consist of roughly 1000 species ^1^. These microorganisms are important for the body’s maintenance of health by maintaining oral homeostasis and protecting the oral cavity ^3,4, 5^. Furthermore, they are involved in metabolic, physiological, and immunological functions, some of which include food digestion and nutrition, metabolic regulation, detoxification of environmental chemicals, prevention of disease-promoting microorganisms, and development and regulation of the immune system to maintain the balance of pro-inflammatory and anti-inflammatory processes ^2^.

When the homeostasis of the microbial communities is no longer intact, the opportunity for diseases can arise. Some of those oral diseases include periodontitis ^6–11^, gingivitis ^7,12,13^, dental caries ^14,15^, oral cancer ^16,17^, and esophageal cancer ^18^. The subgingival pocket provides an ideal environment for the growth of opportunistic anaerobic microorganisms that can more easily translocate into the bloodstream and potentially cause systemic problems. For example, periodontitis is associated with the red complex that appears in the later stages of biofilm development. It comprises species that are considered periodontal pathobionts *(Porphyromonas gingivalis, Treponema denticola*, and *Tannerella forsythia*) that can lead to the development of periodontitis ^19^.

The presence of certain microorganisms in the subgingival pockets can also lead to systemic diseases and illnesses such as colorectal cancer, pancreatic cancer, cystic fibrosis, cardiovascular disease, rheumatoid arthritis, Alzheimer’s disease, and diabetes ^18,20^. For example, *P. gingivalis* and *P. intermedia* form biofilm structures in the subgingival pocket that disrupt the connective tissue, periodontal ligaments, and bones, allowing pathogenic microorganisms access to the bloodstream where atherosclerotic plaques are then formed ^21^. In addition, bacteria can interact with, cause, and elicit responses from the immune system. This is accomplished by the binding of bacteria to the epithelial and endothelial cells, where they can be transported to other locations of the body to begin colonization and evasion of the host defense ^22^. Prolonged inflammation can then lead to chronic inflammatory and systemic diseases, negatively impacting host health ^18^.

Since the subgingival pocket is an ideal environment for these pathobionts, subgingival samples are commonly collected, also known as subgingival plaque samples. The methods of collection include a curette, which has been shown to be difficult to use and requires a dental professional ^23–25^. Another method of collection is a paper point, which is difficult to do alone, and the fragile nature of the paper makes sampling difficult and imprecise ^26^.

The most common method used to analyze the oral microbiome has been 16S rRNA gene sequencing and PCR ^24,25,27,28^. There are a few studies that have metagenomically or metatranscriptomically analyzed subgingival samples from participants who had oral diseases, such as gingivitis or periodontitis ^23,29,30^. Metatranscriptomics (unbiased RNA sequencing) reveals the behavior of the microbiome by quantifying the gene expression profiles of the microbial community ^31^. This technology is important to determine if microorganisms are commensal or dysbiotic, in order to identify the presence of disease and its progression. With this information, we can improve our understanding of how individual bacteria and microbial communities function in human health and diseases. Metatranscriptomics has recently been shown to reveal more relevant microbiome features (their gene expression) than can be measured by traditional DNA profiling methods (16S and metagenomics) that only capture compositional data ^32^.

Herein we validate a very easy, non-invasive, and self-administered method of subgingival sample collection (subgingival-P) that could replace the curette or paper point collection while still obtaining similar quality of microbiome taxonomic and gene expression data. Subgingival samples were collected using a paper point to compare with subgingival-P samples that were collected with a swab along the tooth-gum interface. In our study, we collected subgingival and subgingival-P samples from three teeth of 20 study participants to determine the metatranscriptomic similarities between the two sampling methods. We performed a metatranscriptomic analysis that is clinically validated, automated and enables further studies on the role of subgingival microbiome in human health.

## Materials and methods

### Ethics statement

All procedures involving human subjects were performed in accordance with the United States ethical standards and approved by a federally-accredited Institutional Review Board (IRB).

Informed consent was obtained from the participants, all of whom were at least 18 years old and resided in the USA at the time. The trial registration number on ClinicalTrials.gov is NCT05672953.

### Sample collection

Samples were collected from 20 participants, three of whom used mouthwash regularly (Donor 5, Donor 6, and Donor 7). PerioPaper strips (Oraflow Inc. #593520) were used to collect subgingival samples. The subgingival sample was collected by inserting the PerioPaper strip into the pocket between the tooth and gum and held in place for 30 seconds (Figure 1A). Subgingival samples were collected for tooth 5, tooth 8, and tooth 12 (Table S1). After collection, the PerioPaper strips were placed into a 2 mL tube containing 1 mL of RNA Preservation Buffer (RPB), which was previously validated to preserve RNA for up to 28 days at ambient temperatures ^33, 34,35^. The subgingival-P samples were collected from the same teeth using a flocked swab (Puritan 25-3206-H 20MM). The subgingival-P samples were collected by horizontally placing the flocked swab at the tooth-gum interface, then rotating the swab into the gums while gently pushing for 15 seconds (Figure 1B). After collection, swab tips were placed into a bead beating tube containing RPB and broken off.

**Figure 1:**
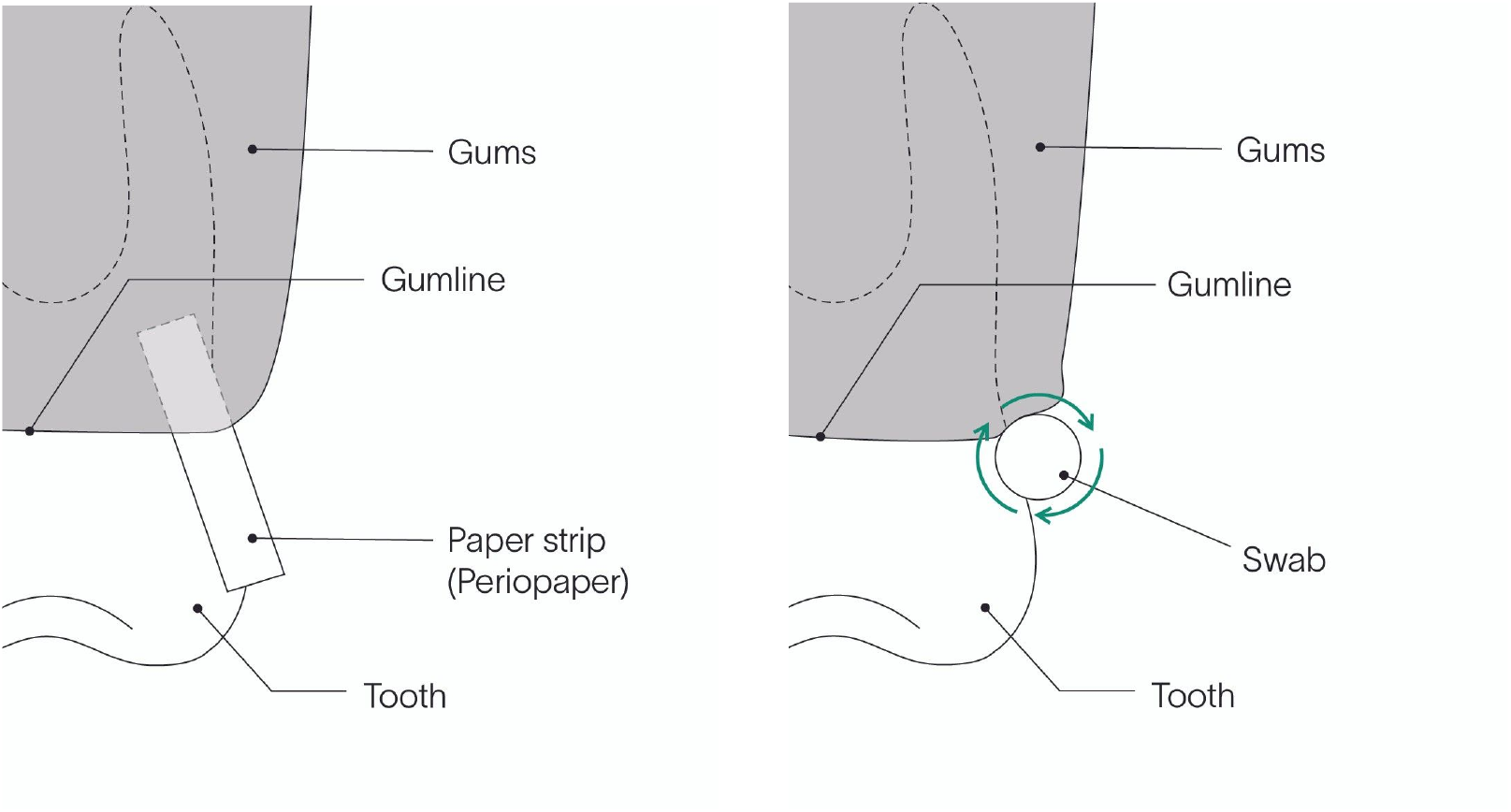
(A) Subgingival sample collection with Periopaper. The PerioPaper strip was inserted into the pocket between the tooth and gum and held in place for 30 seconds. (B) subgingival-P sample collection with swab. A flocked swab was placed horizontally at the tooth-gum interface. The swab was then rotated while gently pushing toward the gums for 15 seconds.

### Laboratory methods

All samples were analyzed in a CLIA laboratory using clinically validated methods. Samples were processed by liquid handlers in a 96-well format. Each batch of 96 samples included external positive and negative process controls, to ensure acceptable method performance. The samples underwent mechanical and chemical lysis and library preparation as previously described ^35^. Briefly, the RNA was extracted using silica beads and a series of washes, followed by elution in molecular biology grade water. DNA was degraded using RNase-free DNase.

Prokaryotic and human rRNAs were removed using a subtractive hybridization method: biotinylated DNA probes with sequences complementary to microbial and human rRNAs were added to total RNA, the mixture was heated and cooled, and the probe-rRNA complexes were removed using magnetic streptavidin beads. The remaining RNAs that contain all other human and microbial RNAs (coding and non-coding) were converted to directional sequencing libraries using unique dual-barcoded adapters and ultrapure reagents. Libraries were pooled and assessed for quality with dsDNA Qubit (Thermo Fisher Scientific) and Fragment Analyzer (Advanced Analytical) methods. Library pools were sequenced on Illumina NovaSeq 6000 instrument using 300 cycle kits.

### Quality control

The quality control includes per-sample and per-batch metrics, such as the level of barcode hopping, batch contamination, positive and negative process controls, DNase efficacy, microbial richness, and number of useful reads obtained per sample. Samples analyzed had to pass quality control acceptance criteria. All other criteria had to pass the laboratory’s standard CLIA quality acceptance criteria, but these two were adjusted for this study: 1. a species richness greater than or equal to 100 and 2. at least 250,000 read pairs aligned to the Viome microbial database.

Samples that did not meet these QC criteria were excluded from data analysis. The samples were split into three groups for comparison. Group 1 compared all subgingival samples for all teeth, a total of 39 samples. Group 2 compared all subgingival-P samples for all teeth, a total of 45 samples. Group 3 compared the subgingival samples and subgingival-P samples for each tooth within each donor, a total of 35 sample pairs (for example, donor 1 subgingival sample tooth 5 was compared to donor 1 subgingival-P sample tooth 5).

### Bioinformatic methods

Viome’s bioinformatics methodologies included quality control, taxonomic classification (at the strain, species, and genus ranks), and microbial quantitative gene expression analysis. Following the quality control, the paired-end reads are aligned to a catalog containing ribosomal RNA (rRNA) and 37,254 genomes spanning archaea, bacteria, fungi, protozoa, phages, and viruses.

Reads that map to rRNA are discarded. Strain-level relative activities were computed from mapped reads via the expectation-maximization (EM) algorithm ^36^. Microbial gene expression levels were computed by mapping paired-end reads to a catalog of 98,527,909 microbial genes and quantifying them with the EM algorithm. Gene-level activities were aggregated into KEGG Orthologs (KOs) ^37^. The identified and quantified active microbial species and KOs for each sample were then used for downstream analyses.

### Read count normalization

Since read counts are compositional, they were transformed prior to calculating any correlations or performing statistical tests on them. Read counts were converted to relative abundance by dividing each read count by the total number of mapped reads in that sample. Multiplicative replacement was then used to replace zeros with a small positive value ensuring that the total for relative abundance still adds up to 1. Finally the centered log ratio (CLR) transformation was applied to go from the simplex to real space.

### Data analysis

The analyses were performed using Python 3.9 standard packages, including Pandas, Numpy, Scipy, Sklearn, Seaborn, Scikit-bio, and Matplotlib.

## Results

### Sequencing metrics

Table 1 summarizes the sequencing metrics for the subgingival and subgingival-P samples.

**Table 1:** Sequencing metrics The average and standard deviation for several important methodological and microbial metrics.

| Sequencing Metric | Subgingival Samples (39) Average | Subgingival Samples (39) Standard Deviation | subgingival-P Samples (45) Average | subgingival-P Samples (45) Standard Deviation |
| --- | --- | --- | --- | --- |
| Total Reads | 10,326,827.21 | 3,578,156.51 | 9,951,015.87 | 3,693,243.09 |
| Percent rRNA | 11.76 | 17.26 | 18.38 | 19.27 |
| Species Richness | 330.59 | 77.84 | 353.04 | 82.45 |
| KO Richness | 4,566.77 | 897.00 | 4,426.42 | 803.60 |
| Bacteria Richness | 313.72 | 77.16 | 338.07 | 81.02 |
| Percent Bacteria | 94.68 | 1.68 | 95.66 | 1.49 |
| Eukaryote Richness | 11.97 | 3.51 | 10.07 | 4.05 |
| Percent Eukaryotes | 3.82 | 1.36 | 2.94 | 1.34 |
| Virus Richness | 4.82 | 2.23 | 4.80 | 1.91 |
| Percent Virus | 1.49 | 0.68 | 1.37 | 0.48 |
| Archaea Richness | 0.08 | 0.27 | 0.11 | 0.32 |
| Percent Archaea | 0.02 | 0.06 | 0.03 | 0.08 |
| Table 1: The average and standard deviation for several important methodological and microbial metrics. |  |  |  |  |

### Intra-donor subgingival samples (Group 1)

In order to determine whether the subgingival microbiome differed between teeth within the same donor, the species relative abundance and KO relative abundance for each tooth within a single donor were compared. The Pearson correlation for species relative abundance had an average of 0.75 (Figure 2A). The Spearman correlation for species relative abundance had an average of 0.72 (Figure 2B). The Pearson correlation for KO relative abundance had an average of 0.77 (Figure 2C). The Spearman correlation for KO relative abundance had an average of 0.80 (Figure 2D). Generally, the correlations for species and KO relative abundances were comparable among teeth from the same donor, indicating that the subgingiva is similar among teeth of the same donor.

**Figure 2:**
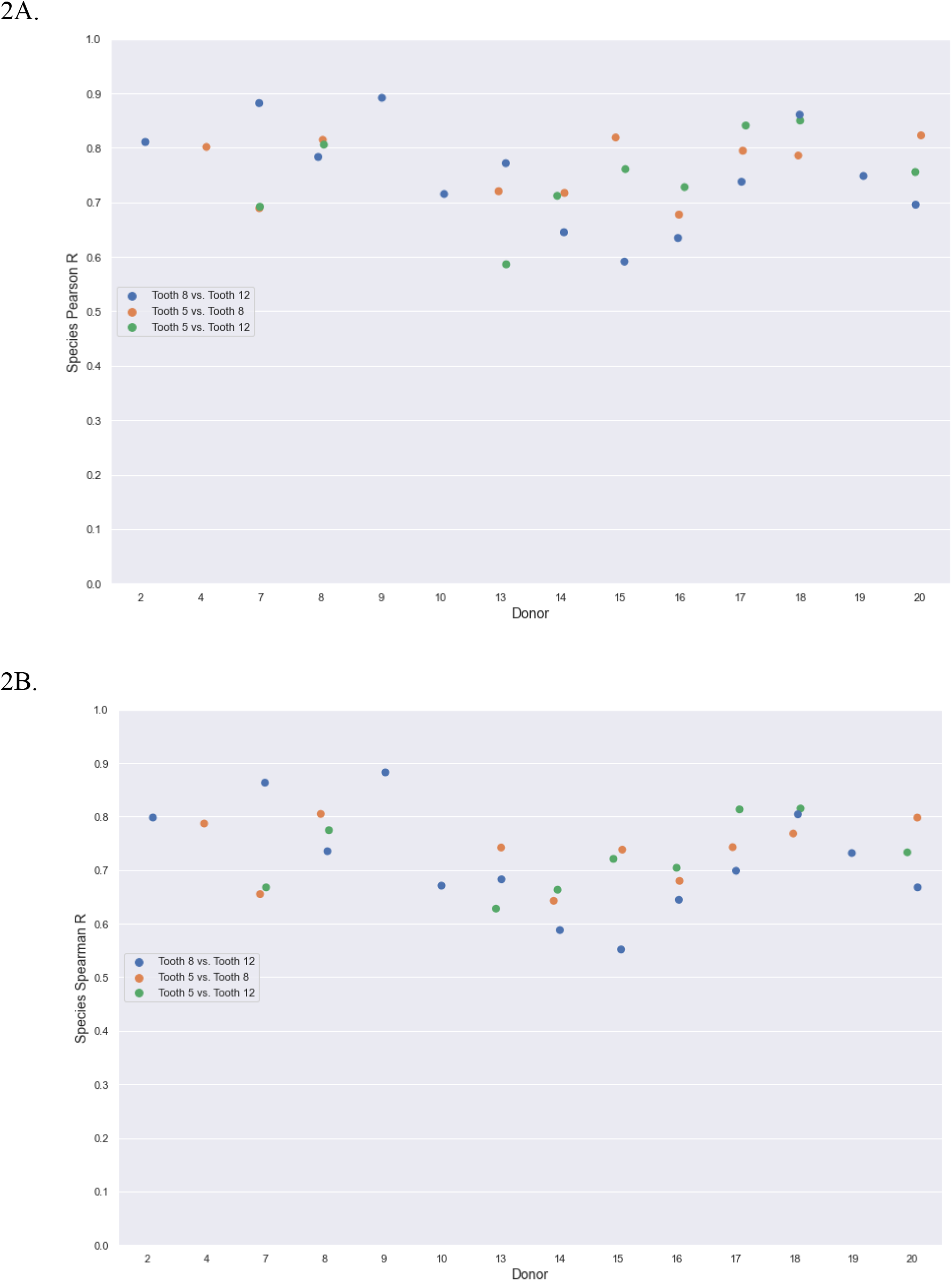
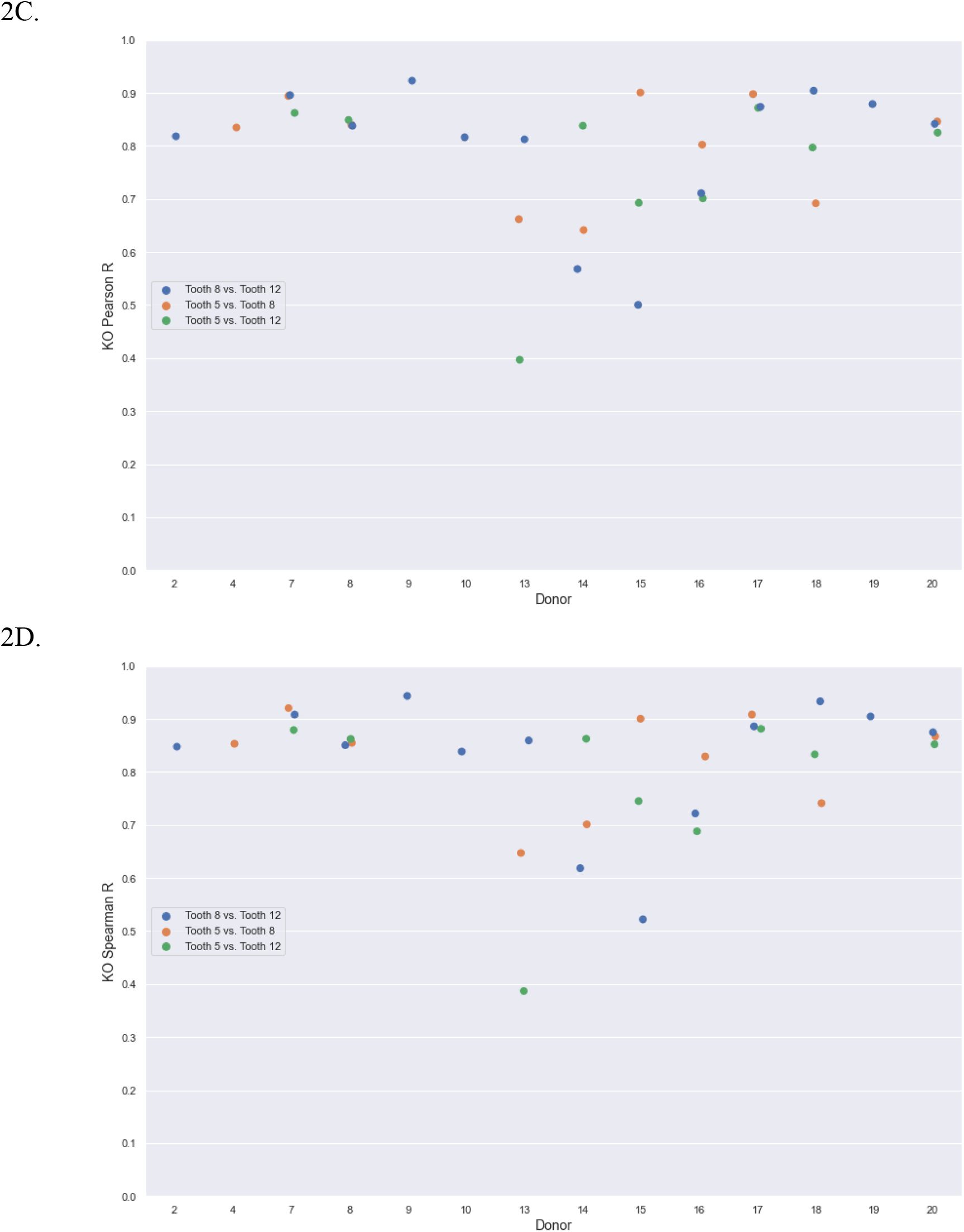
Intra-donor subgingival sample comparisons. A: Pearson correlations of species relative abundance for subgingival samples among the teeth of each donor. B: Spearman correlations of species relative abundance for subgingival samples among the teeth of each donor. C: Pearson correlations of KO relative abundance for subgingival samples among the teeth of each donor. D: Spearman correlations of KO relative abundance for subgingival samples among the teeth of each donor.

### Intra-donor subgingival-P samples (Group 2)

In order to determine whether the subgingival-P microbiome differed among teeth, the species and KO relative abundances for the teeth within a single donor were compared. The Pearson correlation for species relative abundance had an average of 0.81 (Figure 3A). The Spearman correlation for species relative abundance had an average of 0.78 (Figure 3B). The Pearson correlation for KO relative abundance had an average of 0.84 (Figure 3C). The Spearman correlation for KO relative abundance had an average of 0.86 (Figure 3D). Similar to the subgingival sample comparisons, the correlations for species and KO relative abundances were comparable among teeth from the same donor in the subgingival-P samples.

**Figure 3:**
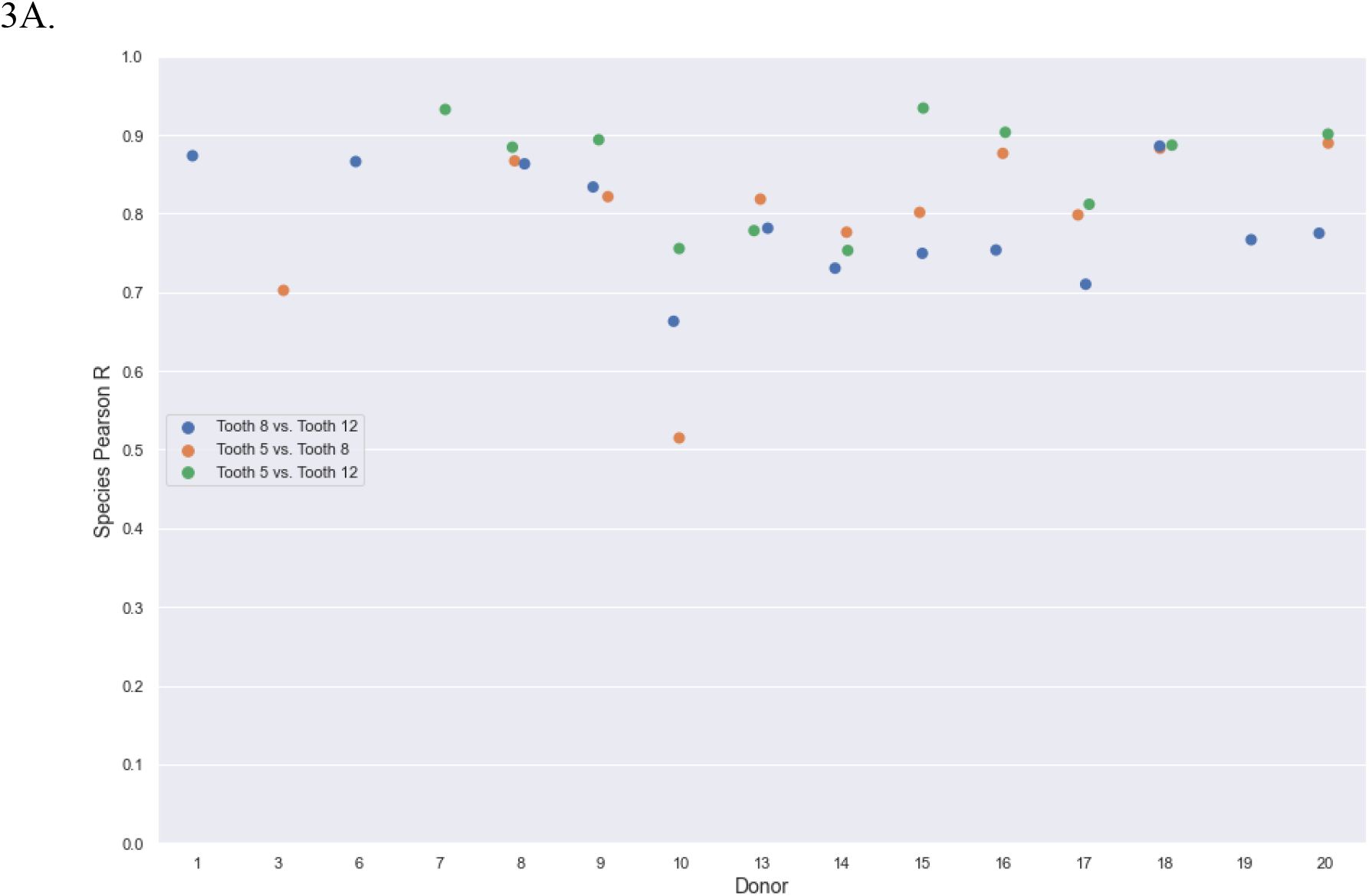
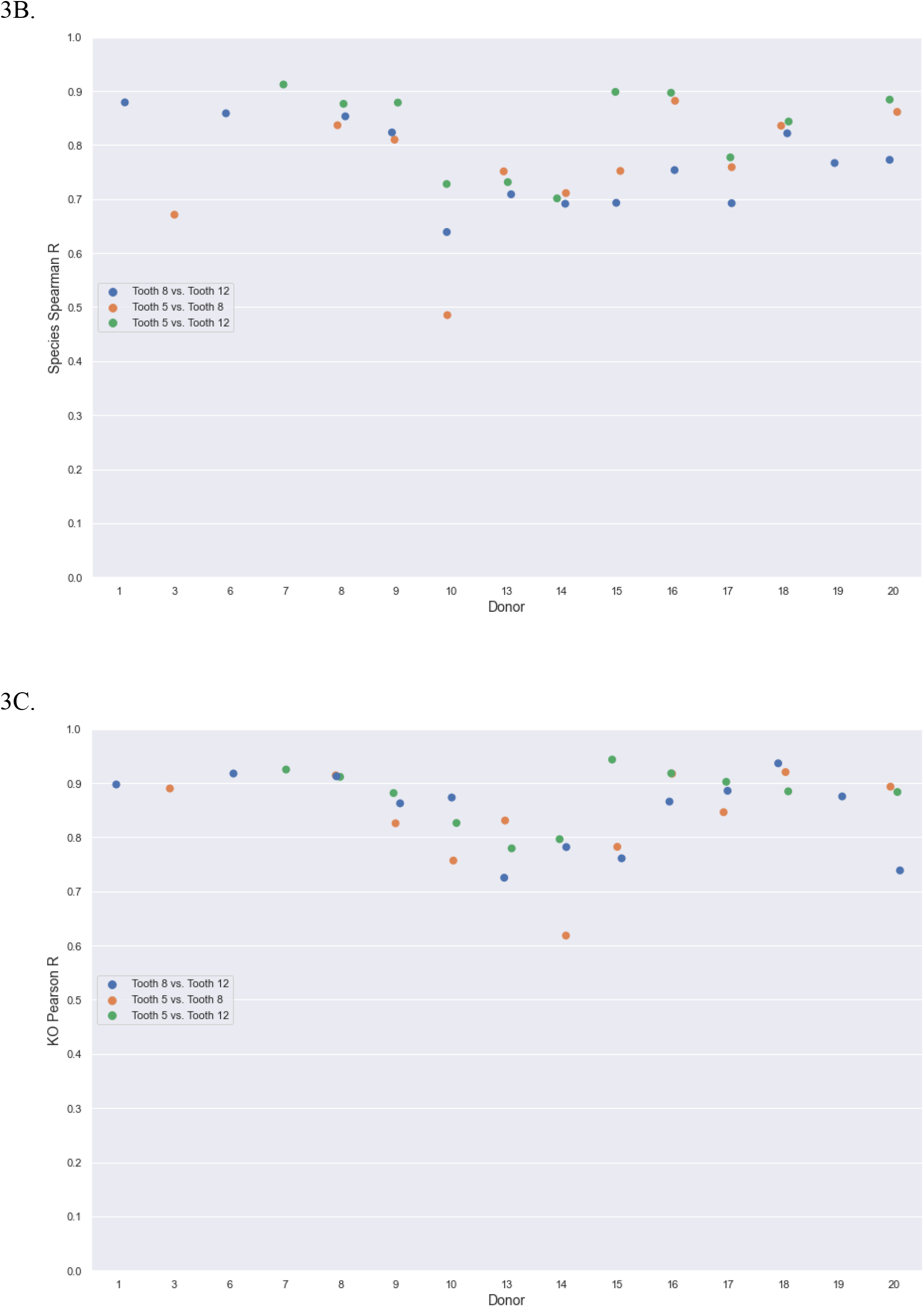
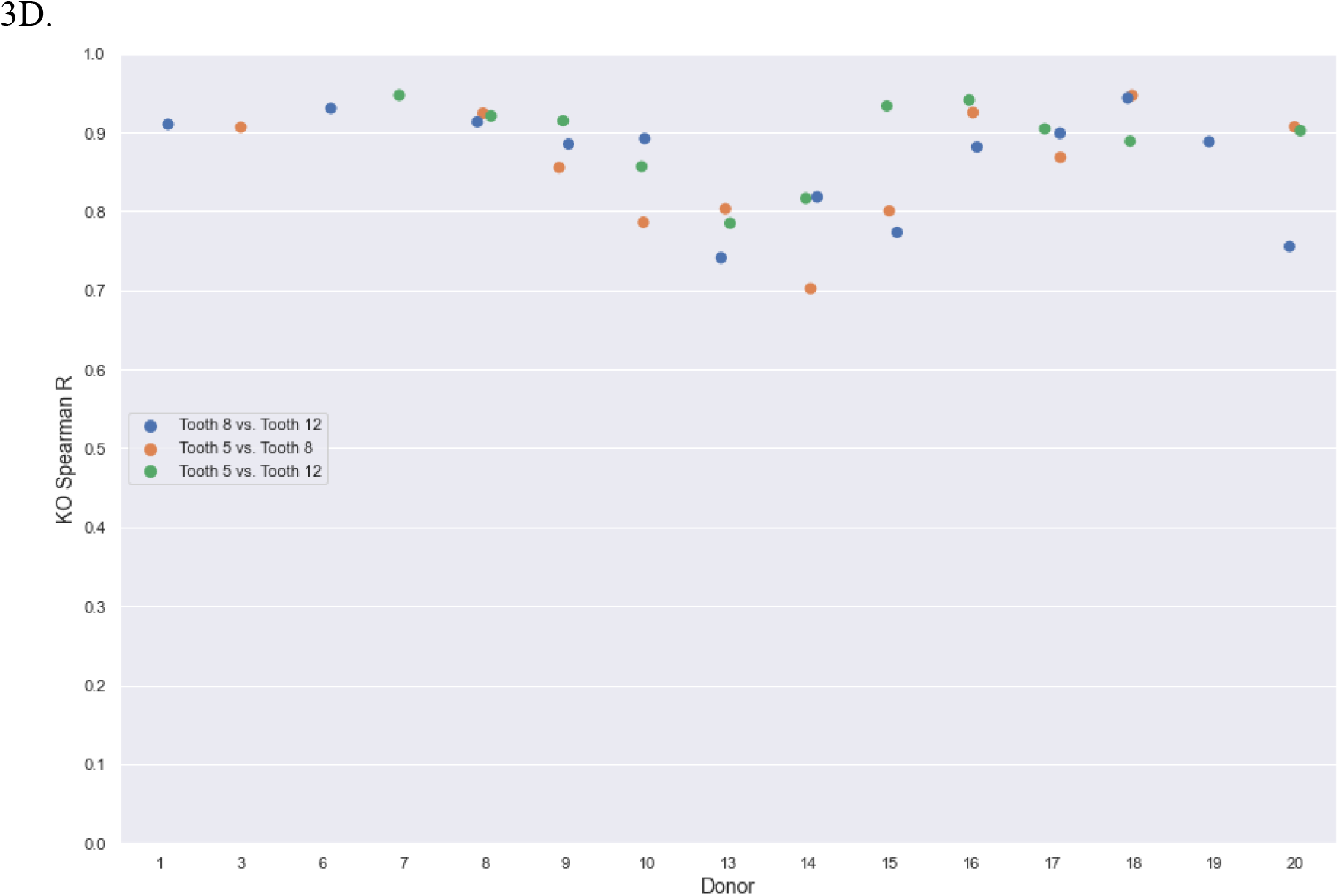
subgingival-P samples vs subgingival-P samples of the same donor. A: Pearson correlations of species relative abundance for subgingival-P samples among the teeth of each donor. B: Spearman correlations of species relative abundance for subgingival-P samples among the teeth of each donor. C: Pearson correlations of KO relative abundance for subgingival-P samples among the teeth of each donor. D: Spearman correlations of KO relative abundance for subgingival-P samples among the teeth of each donor.

### Intra-donor subgingival samples vs subgingival-P samples (Group 3)

In order to determine whether the subgingival microbiome differed from the subgingival-P microbiome, the species and KO relative abundances for the same teeth were compared. The Pearson correlation for species relative abundance had an average of 0.84 (Figure 4A). The Spearman correlation for species relative abundance had an average of 0.82 (Figure 4B). The Pearson correlation for KO relative abundance had an average of 0.81 (Figure 4C). The Spearman correlation for KO relative abundance had an average of 0.82 (Figure 4D). The correlations for species and KO relative abundance were comparable among the subgingival samples and the subgingival-P samples for the same tooth within the same donor.

**Figure 4:**
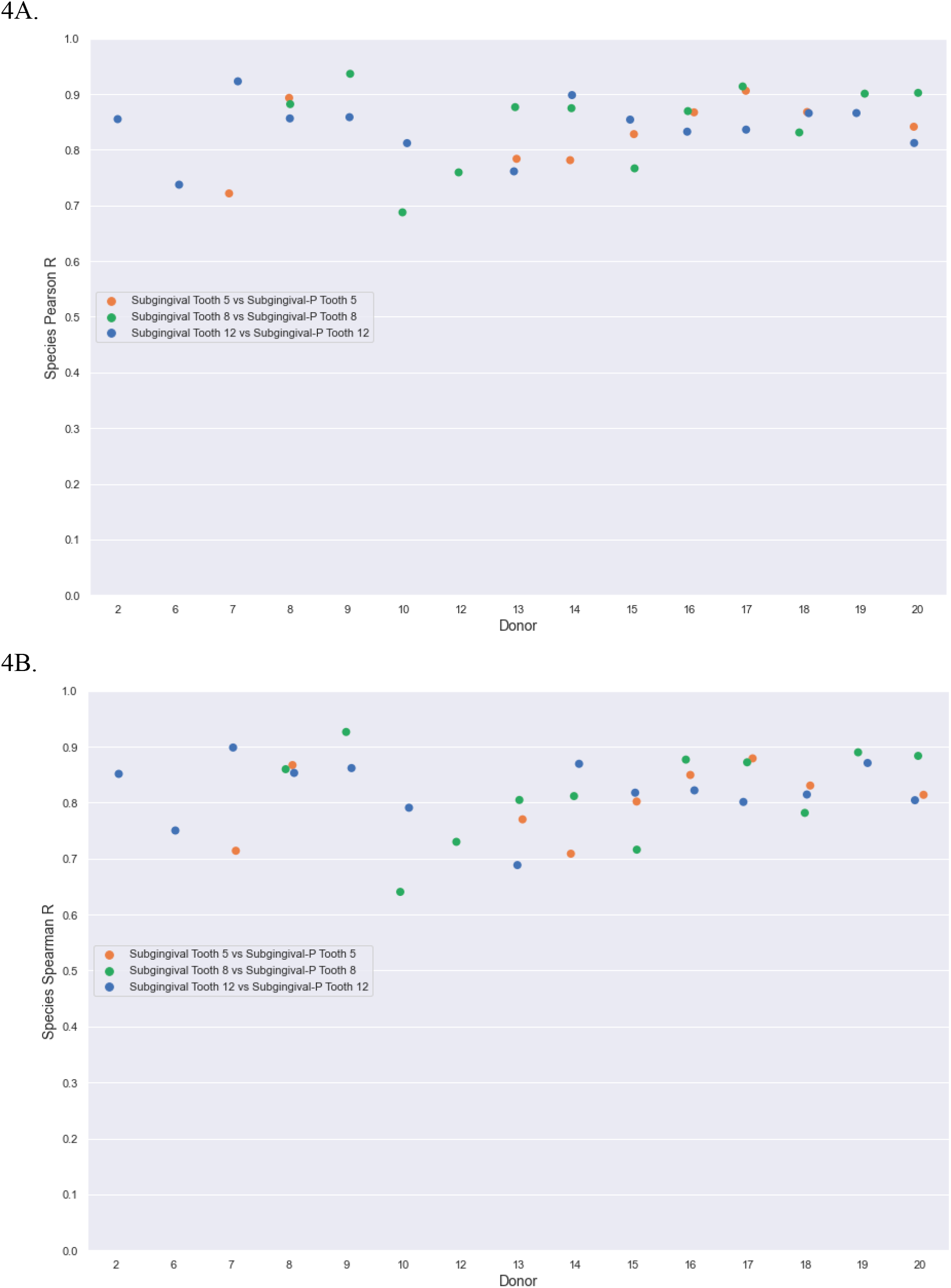
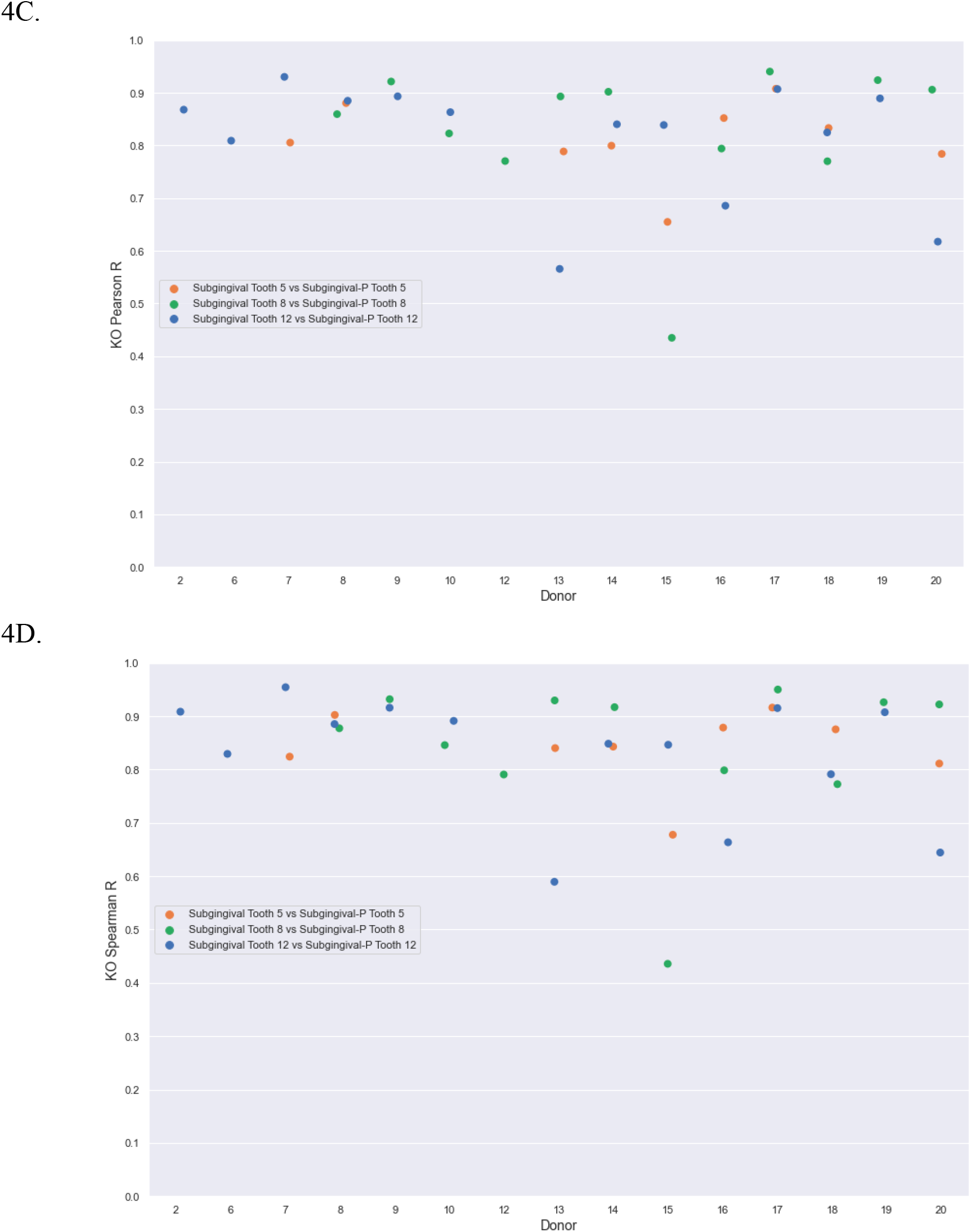
Subgingival samples vs subgingival-P samples. A: Pearson correlations of species relative abundance for subgingival samples vs subgingival-P samples among the teeth of all donors. B: Spearman correlations of species relative abundance for subgingival samples vs subgingival-P samples among the teeth of all donors. C: Pearson correlations of KO relative abundance for subgingival samples vs subgingival-P samples among the teeth of all donors. D: Spearman correlations of KO relative abundance for subgingival samples vs subgingival-P samples among the teeth of all donors.

### Abundance of species unique to subgingival and subgingival-P samples (Group 3)

The species abundance was analyzed to determine the relative abundance and species richness that were unique to the subgingival samples, unique to the subgingival-P samples, and found in both sample locations (overlap). This data analysis was conducted in order to distinguish the ability of the subgingival-P samples to detect the species that are present in the subgingival samples. The average species richness detected in subgingival samples only was 74, the average counts per million (CPM) was 32.11, the standard deviation was 131.8 CPM, and the median was 7.42 CPM. The average species richness detected in subgingival-P samples only was 99, the average was 23.30 CPM, the standard deviation was 109.98 CPM, and the median was 4.54 CPM. The average species richness detected in both the subgingival and subgingival-P samples was 246, the average abundance was 4423.82 CPM, the standard deviation was 29212.6 CPM, and the median was 106.37 CPM. The species relative abundance of the overlap is significantly higher than the relative abundance of species found in subgingival samples or subgingival-P samples only (average p-value <0.05). The overlap (species found in both the subgingival and subgingival-P samples) showed significantly higher species abundance, average species CPM, and median species CPM than those metrics in the subgingival samples only and the subgingival-P samples only. This shows that low abundance species detected in subgingival samples may not always be detected in the corresponding subgingival-P samples, but that the majority of more highly expressed organisms are found in both locations. This is likely due to the sensitivity limit of the method, which can easily be improved with deeper sequencing.

### Correlation of pathobionts in subgingival samples vs subgingival-P samples (Group 3)

The presence and abundance of 21 oral pathobionts (Table S1) were analyzed between subgingival and subgingival-P samples. The pathobionts that were chosen for analysis have been implicated in oral and systemic diseases. The Pearson correlation for species relative abundance had an average of 0.9 (Figure 6A). The Spearman correlation for species relative abundance had an average of 0.88 (Figure 6B). The correlations of the species abundance of the 21 pathobionts between the subgingival and subgingival-P samples for all donors was very high.

**Figure 5:**
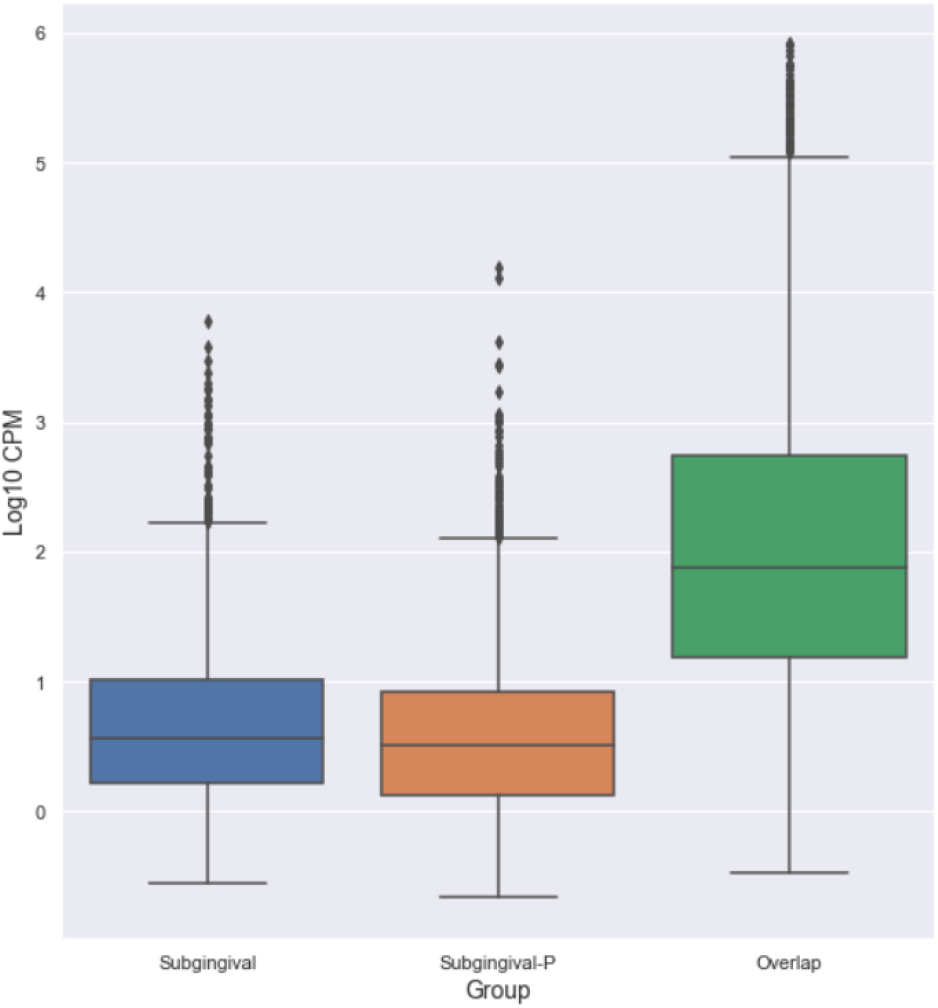
Abundance of unique and overlapping species in subgingival and subgingival-P samples. Species log10 Counts Per Million (CPM) for species detected in subgingival samples only, for species detected in subgingival-P samples only, and for species detected in both subgingival and subgingival-P samples.

**Figure 6:**
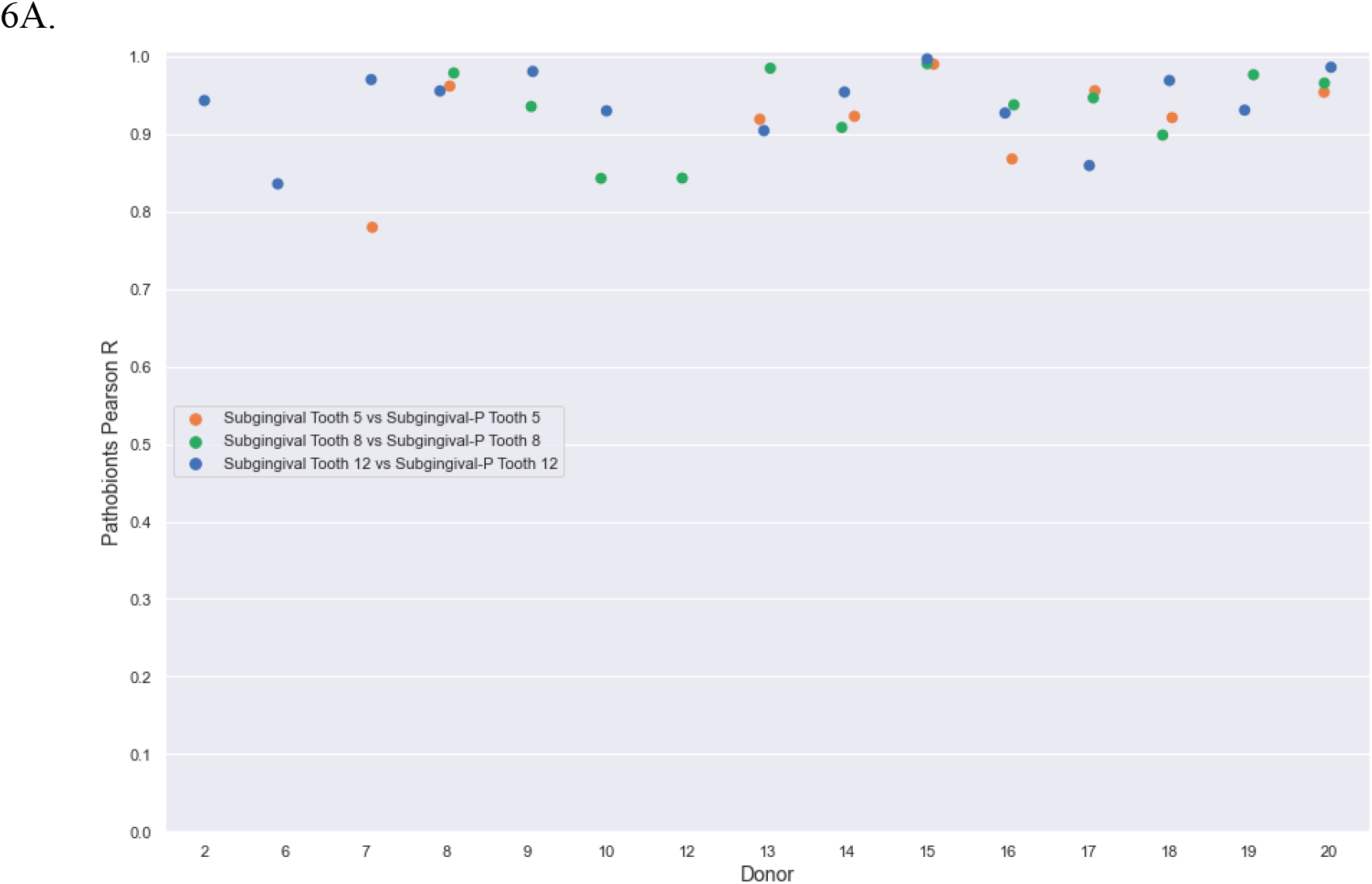
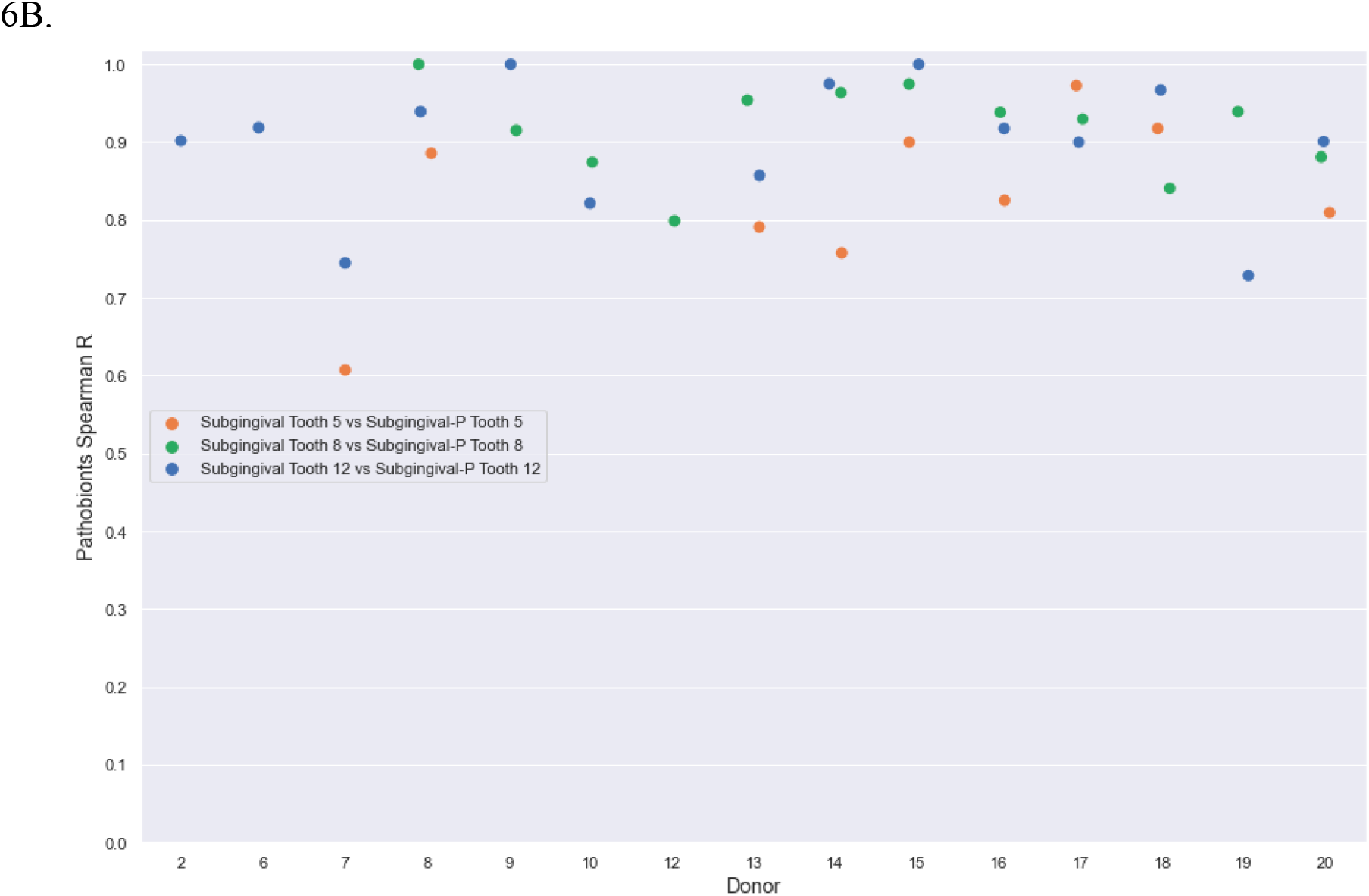
Species abundance correlations of 21 pathobionts between subgingival and subgingival-P samples. A: Pearson correlations of pathobionts relative abundance for subgingival samples vs subgingival-P samples within each tooth. B: Spearman correlations of pathobionts relative abundance for subgingival samples vs subgingival-P samples within each tooth.

### Prevalence of pathobionts in subgingival samples vs subgingival-P samples (group 3)

The ability of the subgingival-P samples to detect the pathobionts (list provided in Table S1) compared to the subgingival samples was assessed to determine if the subgingival-P sampling method compared well to the subgingival method for the detection of pathobionts. The percentage of cases where pathobionts were present in the subgingival-P samples when they were detected in the subgingival samples for each donor’s tooth was calculated (Figure 7).

**Figure 7:**
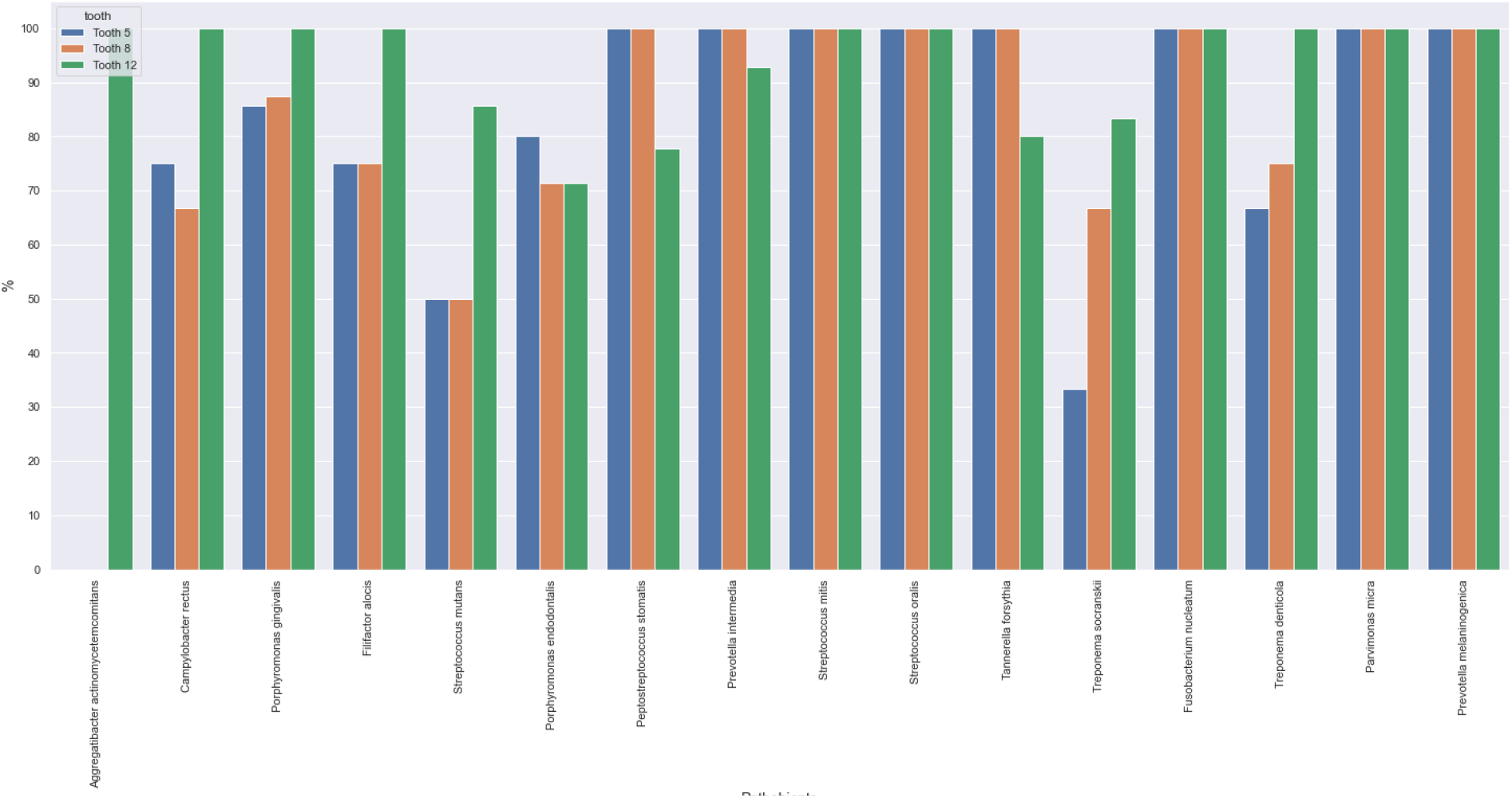
Percentage of cases in which the pathobionts that were detected in subgingival samples were also detected in subgingival-P samples for each tooth of all donors.

## Discussion

The subgingival pocket is sampled in order to identify the microorganisms that are involved in the initiation and progression of certain oral and systemic diseases, such as periodontitis, gingivitis, arthritis, and Alzheimer’s disease. The most common methods of sample collection are curettes and paper points. Both of these have significant drawbacks: curettes require professionals and can be painful, while paper points are flimsy and difficult to use. These drawbacks can impact sample collection and data integrity. In addition, it limits studies to healthcare provider facilities, thus limiting the diversity of the study participants and enrollment rates for clinical studies. For these reasons, a simpler method of collection would be beneficial. We developed a method of collecting subgingival samples by rotating a surface swab at the gumline and named it subgingival-P sampling. Using a metatranscriptomic analysis, we show that subgingival-P samples can identify and quantify the microorganisms found in the subgingival pocket. Importantly, subgingival-P samples can easily be collected by anyone, anywhere.

A critical aspect of any clinically useful sampling method is to ensure that the heterogeneity of the sampling sites is not so large as to make it difficult to identify biomarkers. To determine the innate variability of the microbial composition and functions in the subgingiva, the subgingival samples and subgingival-P samples were collected on three different teeth of 20 donors. The correlations for both gene expression and the relative abundance of microorganisms were high between biological replicates for each sample type. Therefore, the subgingival and subgingival-P sampling shows high homogeneity among teeth, indicating that sampling different teeth within the same person will yield comparable results. While we did observe high correlation between the species abundance among the subgingival samples and the subgingival-P samples, there was some variability in KO abundance comparisons. The gene expression of the microorganisms differed slightly between the teeth for both sample types. Interestingly, a previous study found that the microbial community structures and functions in healthy teeth of an individual can differ from teeth that are progressing to gingivitis ^29^. While we did not explicitly assess oral disease states of the participants, it is possible that the larger differences in KO correlations between teeth for some donors could reflect the progression of some teeth into a diseased state. Therefore, an individual that is being tested for an oral disease may need to get multiple teeth sampled to ensure identification of the functions of certain disease states.

For each tooth sampled, the subgingival and subgingival-P collections were compared for species abundance and KO abundance to ensure that the results between the two sites were comparable. The functions of certain microorganisms can lead to disease progression, therefore, it is vital that gene expression between the subgingival and subgingival-P samples is preserved so that this critical information is maintained across sampling sites. The data analysis showed high species and KO correlations. These results demonstrate that the subgingival-P sampling is generally comparable to the subgingival environment in terms of microbial and KO relative abundance. The subgingival-P sampling is therefore an adequate proxy for the subgingiva.

Although the subgingival and the subgingival-P sampling methods are comparable, there were small differences in the species detected. The data analysis showed that some species in the subgingival samples were absent from the subgingival-P samples and vice versa. However, the species unique to each sample collection location made up a small relative abundance (Figure 5). We conclude that the reason for not detecting some species in the subgingival-P samples was simply limited by the depth of sequencing, and generating more sequencing data would minimize, and potentially eliminate these differences.

The detection of pathobionts may be clinically important because they are strongly associated with the onset and progression of oral and systemic diseases, as previously described ^2^. Our data analysis demonstrated that the pathobionts detected in the subgingival samples were also detected in the subgingival-P samples in high percentages (Figure 7). The pathobionts that were not detected in high percentages, such as *Aggregatibacter actinomycetemcomitans, Streptococcus mutans*, and *Treponema socranskii*, can be explained by the low prevalence in the subgingival samples (Figure S2). Therefore, the subgingival-P samples are sufficiently sensitive for the detection of clinically relevant pathobionts. This is likely important for the diagnosis and prevention of oral and systemic diseases. Clinicians in health facilities and individuals at home could collect a subgingival-P sample easily. Once analyzed, the disease state and disease progression could be determined by analyzing the presence and abundance of pathobionts.

This study has a few limitations. We did not control for the health status of the donors; if a donor had healthy or diseased teeth, it was not known. Metatranscriptomic analysis of the subgingival vs subgingival-P samples on participants with periodontitis or gingivitis will be very useful in order to measure the effectiveness of detection between the two methods.

Herein we were able to demonstrate that subgingival-P samples are a good proxy for subgingival samples. The subgingival-P samples are comparable to subgingival samples in terms of the detection of microorganisms, their relative abundance, and the quantification of microbial gene expression. Subgingival-P samples are much easier to collect than subgingival samples and can be collected at home by patients or study participants. This increased accessibility to sampling the subgingival microbiome could dramatically increase understanding and prevention of oral health and disease.

## Supplemental information

**Figure S1:**
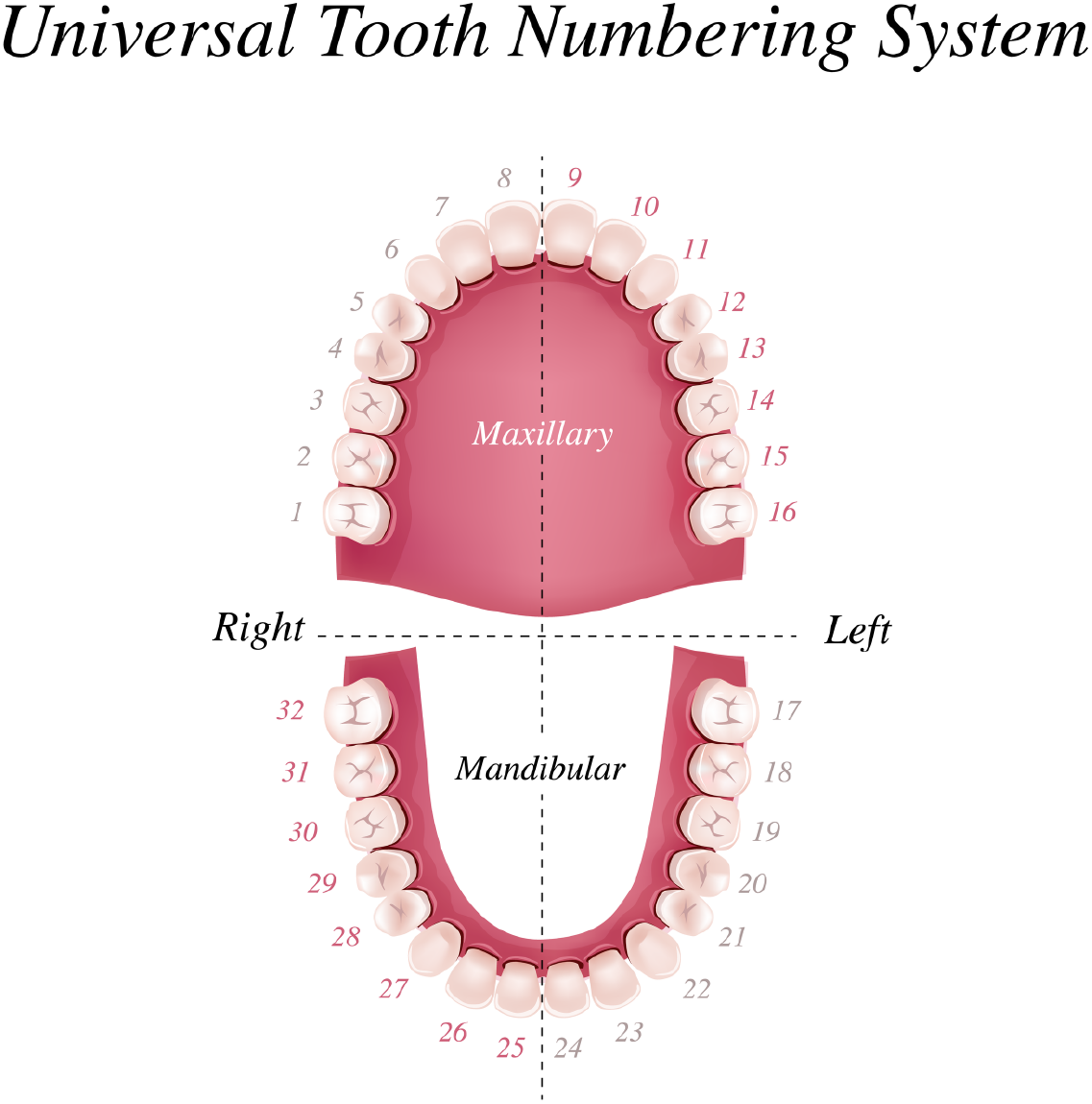
Tooth numbering system. Tooth 5, Tooth 8 and Tooth 12 were sampled

**Table S1:** List of 21 pathobionts The table contains the list of pathobionts investigated in the subgingival and subgingival-P samples. Pathobionts with an asterisk were not detected in the subgingival samples or the subgingival-P samples.

| Pathobiont |
| --- |
| Aggregatibacter actinomycetemcomitans |
| Campylobacter rectus |
| Porphyromonas gingivalis |
| Filifactor alocis |
| Streptococcus mutans |
| Porphyromonas endodontalis |
| Peptostreptococcus stomatis |
| Prevotella intermedia |
| Streptococcus mitis |
| Streptococcus oralis |
| Tannerella forsythia |
| Treponema socranskii |
| Fusobacterium nucleatum |
| Treponema denticola |
| Parvimonas micra |
| Prevotella melaninogenica |
| Methanobrevibacter oralis* |
| Chlamydia pneumoniae* |
| Eubacterium nodatum* |
| Fretibacterium fastidiosum* |
| Cutibacterium acnes (formerly Propionibacterium acnes)* |

### Prevalence of 21 pathobionts in subgingival samples (group 3)

The percentage of cases in which the pathobiont was present in the subgingival sample as well as the subgingival-P samples revealed that some pathobionts were not detected in every single case. To determine if the pathobiont was not detected in the subgingival-P samples because of low presence in the subgingival samples, the percentage of pathobionts present in the subgingival samples was analyzed. The order of the pathobionts follow those of Figure 7.

**Figure S2:**
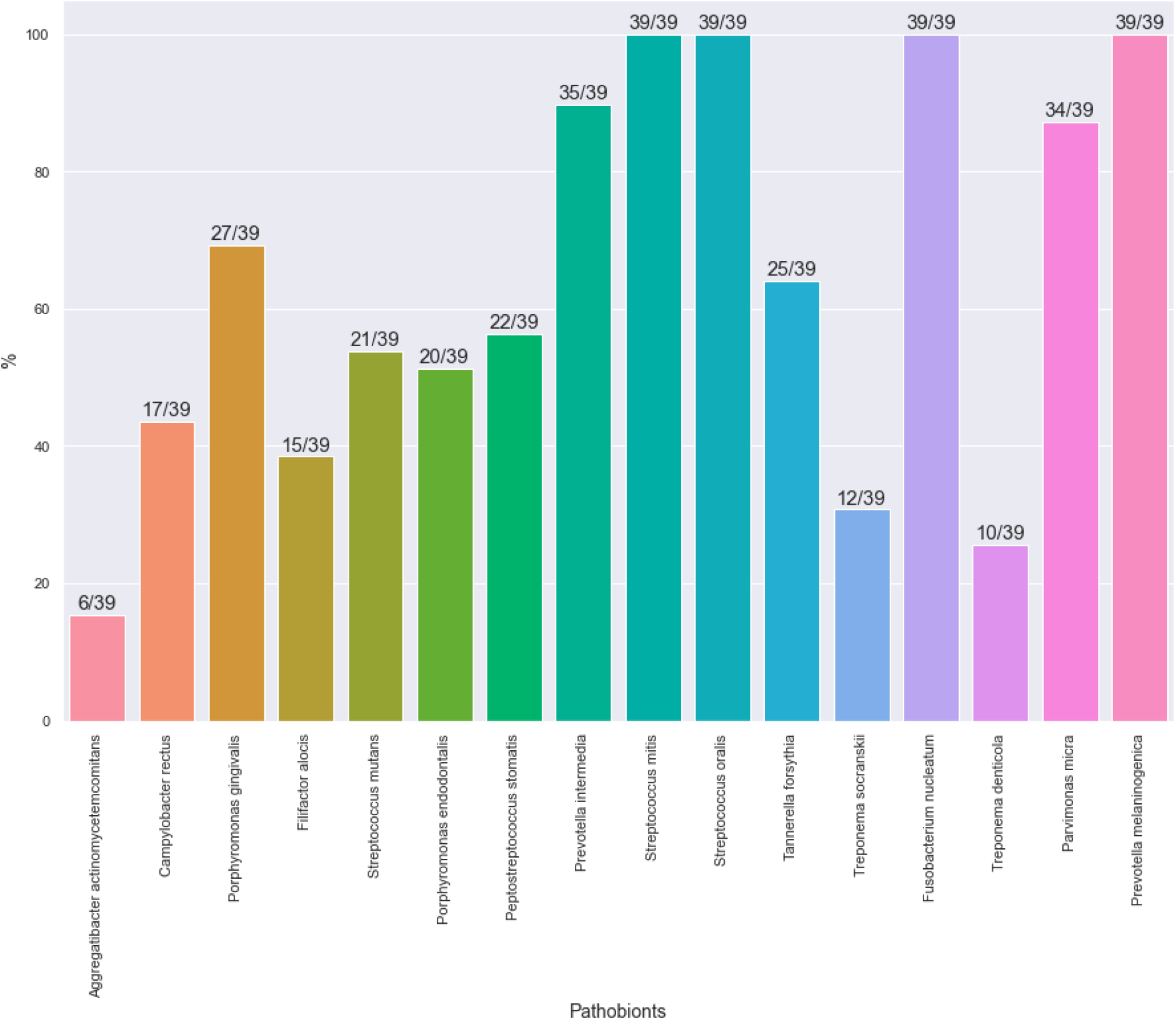
Percentage of subgingival samples in which the pathobionts were detected (for all donors). Only the subgingival samples in group 3 were used for this analysis.

## Data availability

This research was sponsored by Viome and the authors of the paper who have access to the data are employees or scientific collaborators of Viome who have signed contracts with Viome to be bound by Viome’s privacy policy and access restrictions. The sample data and feature matrix for the main discovery cohort (Cohort A) has been made available at figshare, at https://doi.org/10.6084/m9.figshare.13244243. Additional data can be made available through a Data Transfer Agreement that protects the privacy of participants’ data; interested researchers may request at https://www.viome.com/vri/data-access. The information provided by interested researchers on the dataset request form will be used to generate a Statement of Work (no fee SOW) and a Data Transfer Agreement (DTA). The DTA protects the privacy of the participants’ data, and the SOW outlines the planned use of the summary statistics. The SOW and DTA will need to be signed by your institution first, and then Viome, before data can be shared. If you are collaborating with investigators at multiple institutions and those institutions must also receive copies of Viome summary statistics, please have a PI from each institution fill out the form to ensure all parties receive access to datasets within a similar timeframe. Finally, please note that each signed SOW and DTA allows use of Viome data only by the signatory institution and its personnel. Each institution that wishes to access or use Viome data must have a signed SOW and DTA covering their access to Viome data. Once a valid SOW and a valid DTA are signed off, Viome will transfer data to the researcher for use in the research project described in the SOW.

